# Gene expression profiling of The Cancer Genome Atlas supports an inverse association between body mass index (BMI) and major oesophageal tumour subtypes

**DOI:** 10.1101/378778

**Authors:** Inge Seim, Penny L. Jeffery, Lisa K. Chopin

**Affiliations:** Ghrelin Research Group, Translational Research Institute – Institute of Health and Biomedical Innovation, School of Biomedical Sciences, Queensland University of Technology, Brisbane, Queensland, Australia; Comparative and Endocrine Biology Laboratory, Translational Research Institute – Institute of Health and Biomedical Innovation, School of Biomedical Sciences, Queensland University of Technology, Brisbane, Queensland, Australia; Australian Prostate Cancer Research Centre - Queensland, Institute of Health and Biomedical Innovation, Queensland University of Technology, Princess Alexandra Hospital, Translational Research Institute, Brisbane, Queensland, Australia; Integrative Biology Laboratory, College of Life Sciences, Nanjing Normal University, Nanjing, Jiangsu, China

## Abstract

In the last decade the Cancer Genome Atlas (TCGA) program has revealed significant insights into molecular events of dozens of cancers. These data sets are continuously updated, providing an unprecedented resource to the research community. There is now an emerging link between obesity and the development and progression of cancer. In this study we wished to identify genes related to body mass index (BMI) in TCGA datasets. Supporting epidemiological data, our gene expression profiling analyses suggest that oesophageal adenocarcinoma (EAC) can be considered a true obesity-associated cancer subtype, presenting avenues for prevention and treatment.

## Introduction

Obesity is now considered to be an epidemic. Epidemiological data links overweight and obesity, dietary fats, and a sedentary lifestyle with the risk of developing many types of cancer [1], however, mechanistic data has been relatively sparse. In recent years, the development of powerful experimental methods (such as CRISPR [2]) and new technologies (such as RNA-sequencing [3]) have fuelled research aiming to better understand how obesity affects cells and tissues at the whole-body level. Genetic, epidemiological, and experimental data show that a high-fat diet and obesity can mediate tumour progression in diverse cancers [4, 5]. To contribute to this emerging research effort, we here interrogated The Cancer Genome Atlas (TCGA) gene expression data (RNA-sequencing), with the aim of identifying genes whose expression correlates with body mass index (BMI).

## Material and methods

### Datasets

Using the Genomics Data Commons (GDC) data portal [6, 7] we downloaded clinical, exposure (includes BMI), and gene expression (RNA-sequencing) data using the companion GDC Data Transfer Tool from the following 12 The Cancer Genome Atlas (TCGA) datasets with corresponding BMI records: TCGA-BLCA (bladder urothelial carcinoma; *N*=358), TCGA-CESC (cervical squamous cell carcinoma and endocervical adenocarcinoma; *N*=259), TCGA-CHOL (cholangiocarcinoma; *N*=35), TCGA-COAD (colon adenocarcinoma; *N*=231), TCGA-DLBC (lymphoid neoplasm diffuse large B-cell lymphoma; *N*=48), TCGA-ESCA (oesophageal carcinoma; *N*=152), TCGA-KIRP (kidney renal papillary cell carcinoma; *N*=201), TCGA-LIHC (liver hepatocellular carcinoma; *N*=334), TCGA-READ (rectum adenocarcinoma; *N*=72), TCGA-UCEC (uterine corpus endometrial carcinoma; *N*=511), TCGA-UCS (uterine carcinosarcoma; *N*=51), and TCGA-UVM (uveal melanoma; *N*=53). As we interrogated the entire gene set, we employed normalised gene expression values (FPKM-UQ; Fragments Per Kilobase of transcript per Million mapped reads upper quartile). FPKM-UQ is a modified FPKM calculation in which the total protein-coding read count is replaced by the 75^th^ percentile read count value for the sample to scale the reads (see https://docs.gdc.cancer.gov/Data/Bioinformatics_Pipelines/Expression_mRNA_Pipeline/). GDC data is annotated with Ensembl gene IDs. To obtain classical gene symbols and to classify genes into ‘gene types’ (e.g. protein-coding, lincRNA, antisense, and pseudogene), a gene annotation file (GTF; v22; GRCh38.p2) was downloaded from the GENCODE [8] web site (https://www.gencodegenes.org/releases/22.html) and merged with our output data using a custom R script.

### Identification of BMI-related genes

Gene expression data was parsed using custom bash and R scripts, to generate data frames that could be merged with the associated clinical and exposure GDC data files (includes BMI). BMI was calculated as weight (kg) divided by height (m^2^). In each data set, only primary tumour samples were retained, as recommended [7]. Clinical information was manually interrogated to remove obvious outliers (e.g. extreme BMI values in the 132-415 range which could stem from inputting incorrect patient height or the patient has an unspecified subnormal height condition) in the case of TCGA-UZ-A9PU (TCGA-KIRP), TCGA-NH-A8F7 (TCGA-COAD), TCGA-WN-AB4C (TCGA-KIRP), TCGA-SJ-A6ZI (TCGA-UCEC), and TCGA-DD-A4NR (TCGA-LIHC). Additional TCGA-ESCA (oesophageal cancer) clinical information (including histology) was obtained from Supplementary Table 1 of the associated dataset manuscript [9].

To identify body mass index (BMI)-related genes Spearman’s correlation was used for examining the associations between BMI and gene expression values on an ordinal scale [10, 11]. For each dataset, genes with no expression in ≥ 80% of tumours were excluded from analysis [11] and BMI was transformed on a log-scale. A general rule of thumb is that one requires about 30 samples to assume a normal distribution. A sample size of 30 would result in a Spearman correlation (*rho*) cutoff of 0.36, with *rho* decreasing as sample size increases (see [12]). In our analysis we employed a blanket *rho* cutoff of 0.30 and retained genes with a false-discovery-corrected *P*-value less than 0.01 (Bonferroni-corrected to account for a total of 60,483 genes tested). We also compared groups of cancer patients with a normal body mass index value (lean; BMI <25) to overweight (BMI of 25-30) and obese (BMI >30) patients using a non-parametric Mann-Whitney *U*-test (also known as Wilcoxon rank-sum test; corrected for multiple-testing as above). Groups (e.g. cancer subtype) by BMI category (e.g. lean versus high-BMI) were compared using a Pearson Chi-square (χ^2^) test. Scripts are available at GitHub (https://github.com/sciseim/BMI_MS).

### Survival outcome analysis

Survival data was obtained from a recent comprehensive clinical endpoint outcome (OS, overall survival; PFI, progression-free interval; DFI, disease-free interval; DSS, disease-specific survival) curation and analysis effort of 33 TCGA-datasets [7]. Kaplan-Meier survival analysis [13] was performed with the R package ‘survival’ [14], fitting survival curves (*survfit*) and computing log-rank *P*-values using the *survdiff* function, with *rho*=0 (equivalent to the method employed by UCSC Xena; see https://goo.gl/4knf62). Survival curves were plotted when survival was significantly different between two groups (log-rank *P* ≤ 0.05). Datasets with fewer than 10 events per group can be considered unreliable [7].

## Results and discussion

As of May 2018, The Cancer Genome Atlas (TCGA) program had amassed clinical and molecular data of more than 30 different cancer types. Of these, 12 cancer types had associated body mass index (BMI) data (Table 1). We employed Spearman’s correlation test to assess the correlation between gene expression (RNA-sequencing data) of protein- and non-protein coding genes and BMI (cut-off set at an absolute *rho* of 0.30 and a multiple testing adjusted *P*-value, *Q*, less than 0.01). Using this threshold, we observed significant correlation between gene expression and BMI in four TCGA datasets (summarised in Table 1): TCGA-CESC (cervical squamous cell carcinoma and endocervical adenocarcinoma; 1 gene with negative correlation), TCGA-LIHC (liver hepatocellular carcinoma; 4 positive and 3 negative), TCGA-UCEC (uterine corpus endometrial carcinoma; 6 positive), and TCGA-ESCA (oesophageal carcinoma; 771 positive and 583 negative) (Table 1). Please see Table 2 and S1-S3 Tables for details.

**Table 1.** BMI-related genes identified by Spearman’s correlation test. Abbreviations: BMI, body mass index; *N*_lean_, the number of patients with a BMI below 25; *N*_overweight_, the number of patients with a BMI between 25 and 30; *N*_overweight_, the number of patients with a BMI above 30; *N*_combined_, the number of genes identified by Spearman correlation at a *rho* cut-off of 0.30 and a multiple-testing adjusted *P-*value (Bonferroni) less than 0.01; Up, the number of genes with a positive correlation with BMI; Down, the number of genes with a negative correlation with BMI. BMI distribution is shown as mean (lower quartile-upper quartile). The Cancer Genome Atlas (TCGA) datasets are TCGA-BLCA (bladder urothelial carcinoma), TCGA-CESC (cervical squamous cell carcinoma and endocervical adenocarcinoma), TCGA-CHOL (cholangiocarcinoma), TCGA-COAD (colon adenocarcinoma), TCGA-DLBC (lymphoid neoplasm diffuse large B-cell lymphoma), TCGA-ESCA (oesophageal carcinoma), TCGA-KIRP (kidney renal papillary cell carcinoma), TCGA-LIHC (liver hepatocellular carcinoma), TCGA-READ (rectum adenocarcinoma), TCGA-UCEC (uterine corpus endometrial carcinoma), TCGA-UCS (uterine carcinosarcoma), and TCGA-UVM (uveal melanoma).

| Dataset | BMI distribution<br>($\text{kg m}^{-2}$) | $N_{\text{lean}}$ | $N_{\text{overweight}}$ | $N_{\text{obese}}$ | $N_{\text{cor}}$ | | |
| --- | --- | --- | --- | --- | --- | --- | --- |
|  |  |  |  |  | All | Up | Down |
| TCGA-BLCA | 26.99<br>(23.20-29.63) | 149 | 124 | 85 | 0 | 0 | 0 |
| TCGA-CESC | 27.95<br>(22.62-32.41) | 98 | 75 | 86 | 1 | 0 | 1 |
| TCGA-CHOL | 27.99<br>(24.82-30.22) | 10 | 16 | 9 | 0 | 0 | 0 |
| TCGA-COAD | 28.37<br>(23.85-31.83) | 78 | 80 | 73 | 0 | 0 | 0 |
| TCGA-DLBC | 25.91<br>(22.01-28.37) | 25 | 15 | 8 | 0 | 0 | 0 |
| TCGA-<br>ESCA | 25.07<br>(21.30-27.47) | 84 | 46 | 22 | 1294 | 711 | 583 |
| TCGA-<br>KIRP | 28.92<br>(25.26-32.34) | 50 | 86 | 74 | 0 | 0 | 0 |
| TCGA-<br>LIHC | 25.82<br>(21.76-28.66) | 177 | 90 | 67 | 4 | 2 | 2 |
| TCGA-<br>READ | 27.12<br>(24.60-29.15) | 20 | 37 | 15 | 0 | 0 | 0 |
| TCGA-UCEC | 33.43<br>(26.37-38.83) | 95 | 114 | 302 | 3 | 3 | 0 |
| TCGA-UCS | 29.71<br>(23.95-32.86) | 17 | 14 | 20 | 0 | 0 | 0 |
| TCGA-UVM | 28.68<br>(23.32-30.32) | 17 | 22 | 14 | 0 | 0 | 0 |

**Table 2.**
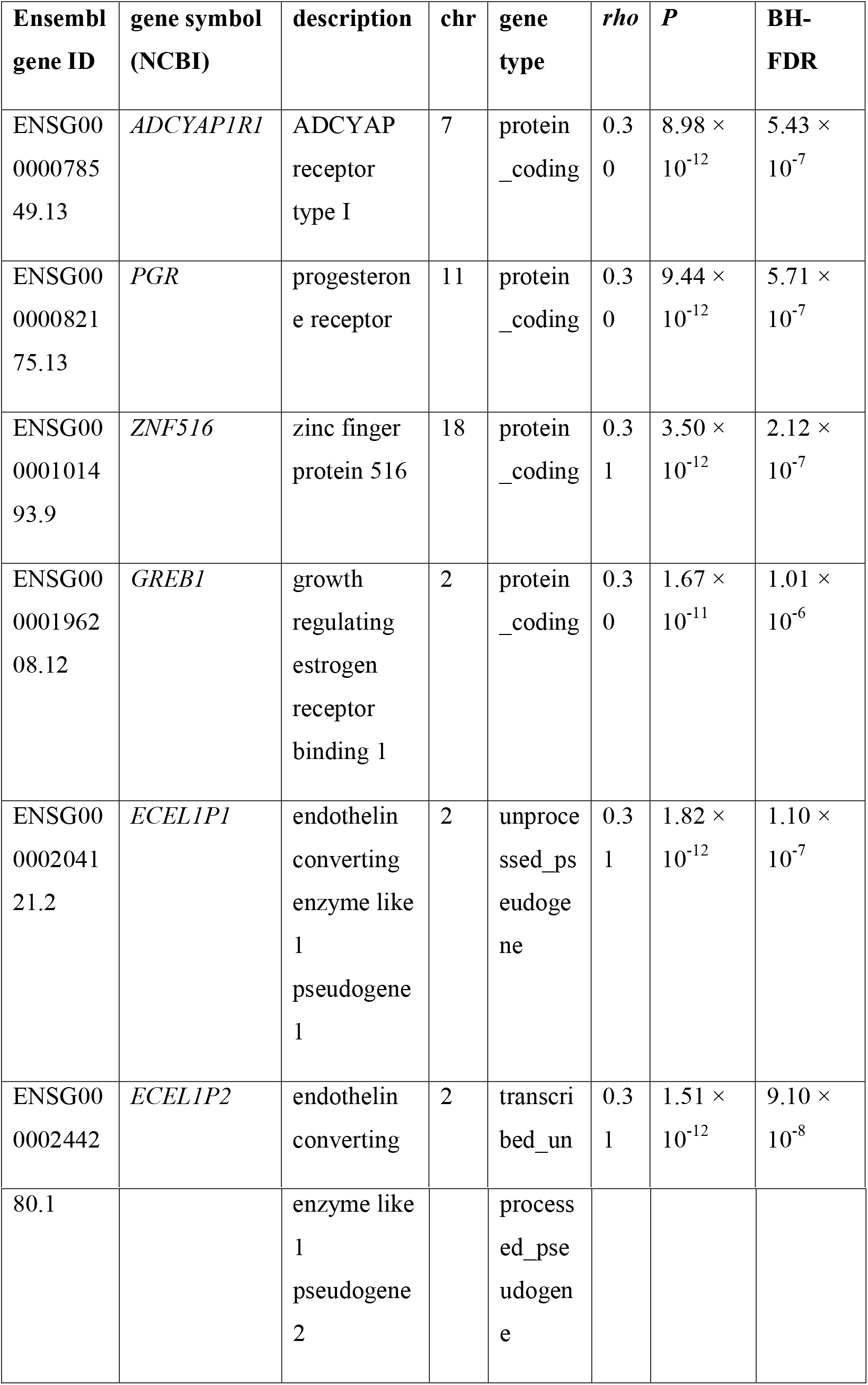
BMI-related genes identified by Spearman correlation test in TCGA-UCEC (uterine corpus endometrial carcinoma). Cutoff set at a Spearman *rho* of 0.30 and a multiple-testing corrected *P*-value less than 0.01 (Bonferroni-adjusted *P, BH-FDR*).

In a previous study which employed an earlier iteration of the TCGA-UCEC endometrial carcinoma dataset, analysis was limited to endometroid tumours and linear regression (with no *r*^2^ cut-off) was performed on protein-coding genes and BMI. Comparing 185 obese (BMI >30) and 105 non-obese patients, 214 genes were associated with body mass index. In our analysis the strength of the relationship between a gene’s expression and BMI was quantified using Spearman correlation analysis. Interrogating a total of 40,669 non-coding and 19,814 protein-coding genes in 302 obese and 213 non-obese TCGA-UCEC patients we identified four protein-coding genes and two non-coding genes with significant associations between expression and BMI. Two protein-coding genes identified are critical regulators of hormone-dependent tumour growth and also identified in the previous study [15]: growth regulating oestrogen receptor binding 1 (*GREB1*) and progesterone receptor (*PGR*) were both positively associated (Spearman *r*=0.30, *Q*=5.71 × 10^-7^ and *r*=0.30, *Q*=1.01 × 10^-6^, respectively) with body mass index (Table 2). A very recent study on an independent clinical data set revealed that PGR expression is indeed correlated with BMI in endometrial cancer [16, 17]. GREB1 is essential for progesterone-driven endometrial decidualisation [18]. Hormone-targeted therapies are a common approach for advanced endometrial cancer and are more effective in patients with progesterone receptor positive tumours [19]. These data suggest that *PGR* and *GREB1* should be further studied to understand the link between endometrial cancer and increasing rates of obesity worldwide [20].

Because a large number of TCGA-ESCA (oesophageal carcinoma) genes (1,294 of a total of 60,483 genes; 2.14%) (Table 1) correlated with BMI, we performed additional analyses on this dataset (see S1 Fig.). First, we repeated the analyses with a stricter Spearman *rho* (0.50) and adjusted *P*-value cut-off (*Q* ≤ 0.001). Under these conditions, 145 genes (0.24%) (S3 Table) were significant. Of these, 86 (59.3%) showed a positive correlation with BMI and 59 (46.7%) a negative correlation. These genes were also significant when BMI was dichotomised into lean and high-BMI (BMI above 25; overweight and obese) (Mann-Whitney *U*-test with *Q* ≤ 0.05). Because GENCODE [8] annotations were employed, BMI-related genes included protein-coding genes (69 with a positive correlation with BMI; 45 with a negative correlation) as well as non-coding RNA genes (e.g. antisense lncRNAs). In an attempt to identify clinico-biologically meaningful groups, we performed unsupervised hierarchical clustering (Euclidian, average) of the expression of the 145 BMI-related genes in the TCGA-ESCA dataset. The resulting heat map revealed a striking difference between the two major subtypes of oesophageal carcinoma, ESCC (squamous cell carcinoma) and EAC (adenocarcinoma) (Fig. 1). EAC tumours largely clustered with high-BMI (overweight and obese). In agreement, stratifying EAC and ESCC into lean (20 EAC and 58 ESCC) and high-BMI (46 EAC and 16 ESCC) groups showed that the subtypes are over-represented in distinct BMI strata (Pearson χ^2^-test, *P*=1.09 × 10^-8^). Fig. 2 illustrates genes with distinct expression in the dataset: Elevated expression of p63 (*TP63*) [9, 21] and the long non-coding RNA *LINC01415* in ESCC; the converse for mucin 13 (*MUC13*) [22] and the long non-coding RNA *LINC00675* [23] in EAC.

**Fig. 1.**
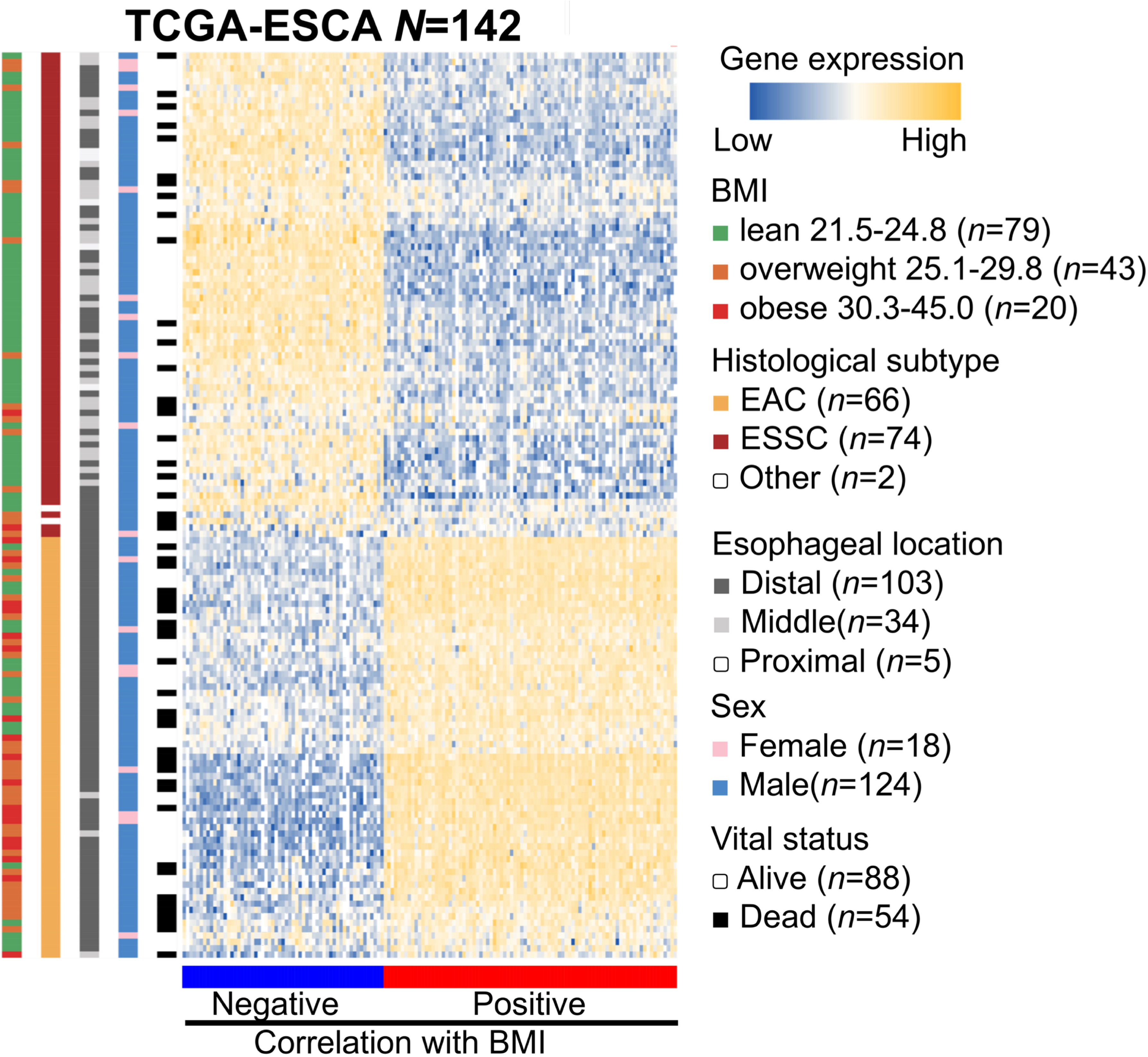
Heat map of gene expression of 145 BMI-related genes in TCGA-ESCA (oesophageal carcinoma). Normalised to depict relative values within rows (samples) with high (yellow) and low expression (blue). Various clinical stratifications are shown as colour bars on the left-hand side; negative (blue) and positive (red) correlation with BMI is shown as a bottom colour bar. Samples with no expression of one or more of the genes were excluded from the heatmap.

**Fig. 2.**
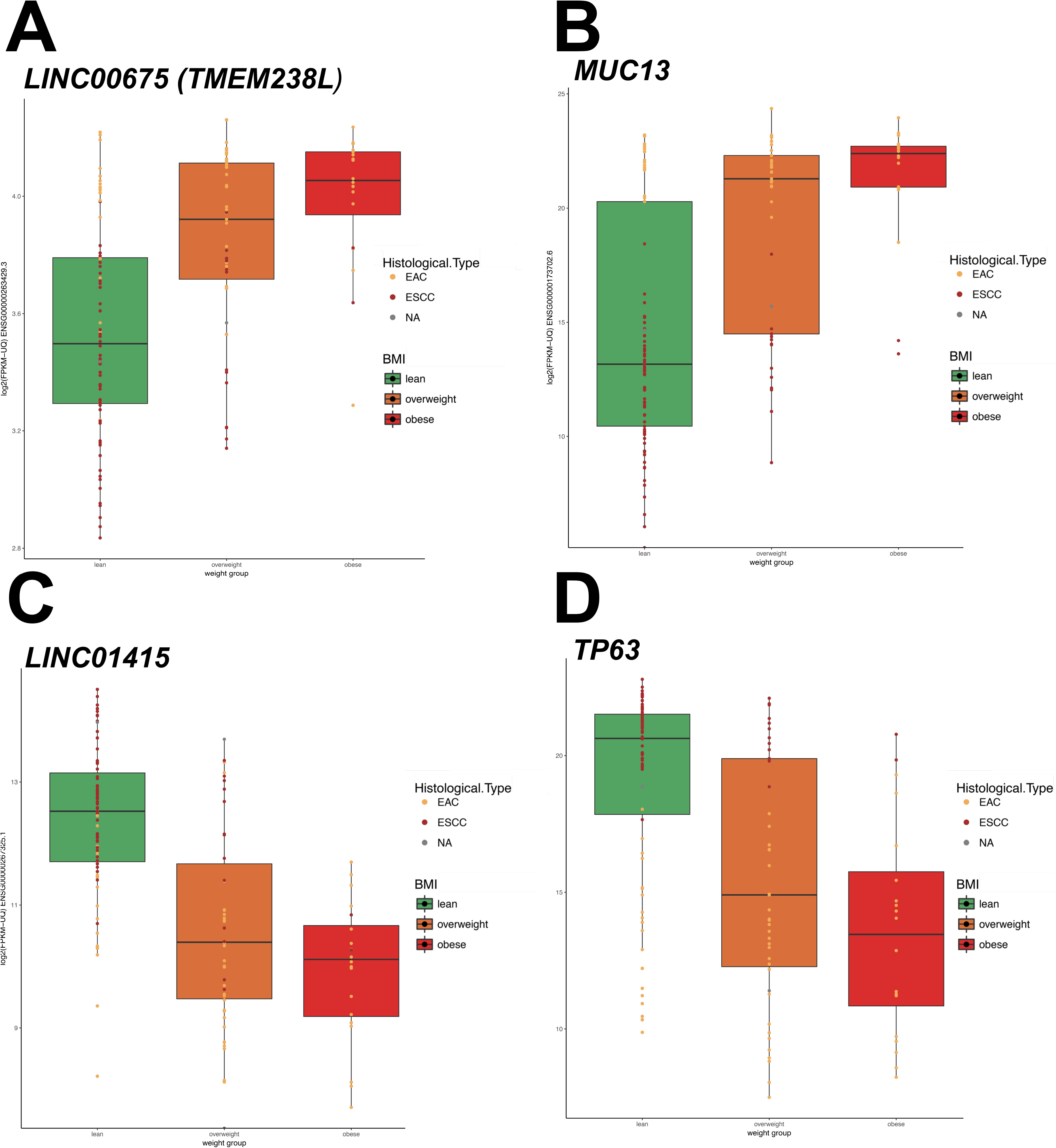
Box plots showing the expression of selected BMI-associated genes in the TCGA-ESCA (oesophageal cancer) data set. The *x*-axis represents samples grouped into lean (BMI <25; in green), overweight (BMI of 25-30; in orange), and obese (BMI >30; in red); the *y*-axis log_2_(FPKM-UQ) gene expression of the gene. The expression of patients with the oesophageal cancer subtypes and EAC (adenocarcinoma) and ESCC (squamous cell carcinoma) are shown as distinctly-coloured dots.

A natural next question to ask is what are the implications from this result? We considered the current literature on the relationship between the oesophagus and BMI, and oesophageal carcinoma and obesity. Recent comparative analysis (employing Spearman correlation) on normal tissues revealed that oesophageal tissue is one of six tissues (the other being adipose, nerve, pancreas, pituitary, and skin) where gene expression is strongly associated with BMI [11]. Oesophageal carcinoma is one of the most aggressive human cancers [24]. The TCGA-ESCA companion manuscript argued that EAC (oesophageal adenocarcinoma) and ESCC (squamous cell carcinoma) should be considered inherently distinct subtypes, developed by lineage-specific alterations [9]. EAC and the gastric tumour subtype chromosomal instability (CIN) are remarkably homogenous, both at the histopathological and molecular level, while ESCC is more similar to squamous cell carcinoma derived from other tissues [9]. Of the two subtypes, ESCC is more prevalent worldwide and high BMI is associated with reduced risk of developing this disease [25, 26]; while EAC is highly prevalent in populations where obesity is common [27]. There has been a shift in the prevalence of EAC in the USA correlating with an increase in obesity and sedentary lifestyle, with EAC increasing from 10% of cases in 1975 to 80% in 2014 [24]. Furthermore, multiple, large population-based (epidemiological studies) studies show a strong correlation between increased BMI and the risk of developing EAC [25, 27, 28]. EAC is prevalent in men, and potentially associated with masculine fat distribution (primarily abdominal) and sex hormones [9, 27]. These epidemiological observations fit well with our data. We also compared the prognosis of the BMI groups using various endpoints, however, there were too few patients with clinical endpoint events when patients were divided on the basis of histological subtypes (S4 Table). Nevertheless, correlating TCGA-ESCA data with BMI identifies a gene set which supports distinct diet-associated origins of oesophageal cancer subtypes.

## Conclusions

In conclusion, by integrating gene expression and BMI data from the TCGA program, we have gained insights into the association between overweight and obesity and cancer. Integration of genomic and clinical datasets should be carefully performed to identify subtypes within each cancer, as recommended by TCGA [7]. Indeed, such efforts are likely to reveal further BMI-associated cancer subtypes. Nevertheless, when distinct subtypes associated with overweight and obesity are evident, we and others [15] have shown that correlating gene expression and BMI alone across an entire data set is informative. We propose that an obesogenic environment is essential for the development of oesophageal adenocarcinoma (EAC), but not oesophageal squamous cell carcinoma (ESCC) – adding further weight to distinct prevention and treatment strategies for these histological subtypes [9]. Future studies, integrating additional genomic datasets (such as epigenome data) and the release of updated clinical information by TCGA is likely to further define the link between a high BMI and cancer.

## Supporting information

**S1 Fig. Overview of body mass index (BMI) distribution in the TCGA-ESCA (oesophageal cancer) data set**. Lean (BMI <25; in green), overweight (BMI of 25-30; in orange), and obese (BMI >30; in red) patients are shown.

**S1 Table. BMI-related genes identified by Spearman correlation test in TCGA-CESC (cervical squamous cell carcinoma and endocervical adenocarcinoma).** Cutoff set at *rho* of 0.30 and a multiple-testing corrected *P*-value less than 0.01 (Bonferroni-adjusted *P*).

**S2 Table. BMI-related genes identified by Spearman correlation test in TCGA-LIHC (liver hepatocellular carcinoma).** Cutoff set at *rho* of 0.30 and a multiple-testing corrected *P*-value less than 0.01 (Bonferroni-adjusted *P*).

**S3 Table. BMI-related genes identified by Spearman correlation test in TCGA-ESCA (esophageal carcinoma).** Cutoff set at *rho* of 0.50 and a multiple-testing corrected P-value less than 0.001 (Bonferroni-adjusted *P*). The following genes have a different symbol in GENCODE: *AFDN-DT* (*MLLT4-AS1*), *PHETA1* (*FAM109A*), and *TMEM238L* (*LINC00675*).

**S4 Table**. **Survival outcome analysis of TCGA-ESCA subsets.** BMI denotes body mass index; EAC. oesophageal adenocarcinoma; ESCC, oesophageal squamous cell carcinoma; *P*, log-rank *P*-value; OS, overall survival; PFI progression-free interval; DFI, disease-free interval; DSS, disease-specific survival.

